## Supplemental materials for "Leveraging synthetic data produced from museum specimens to train adaptable species classification models"

<u>Figs</u>

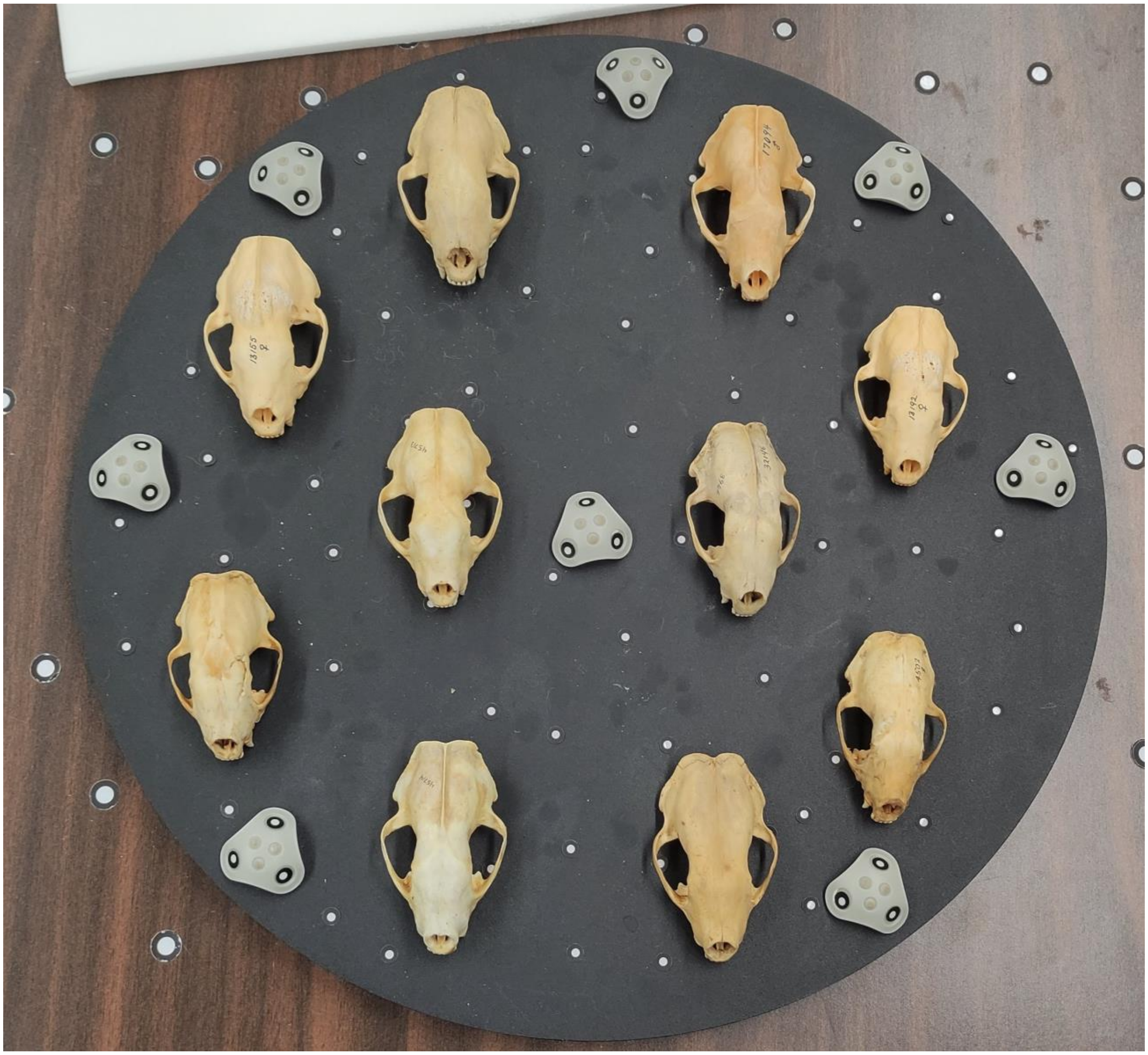

Fig S.1: A photograph of the skull 3D scanning table with 10 skunk (*Mephitis mephitis*) skulls.

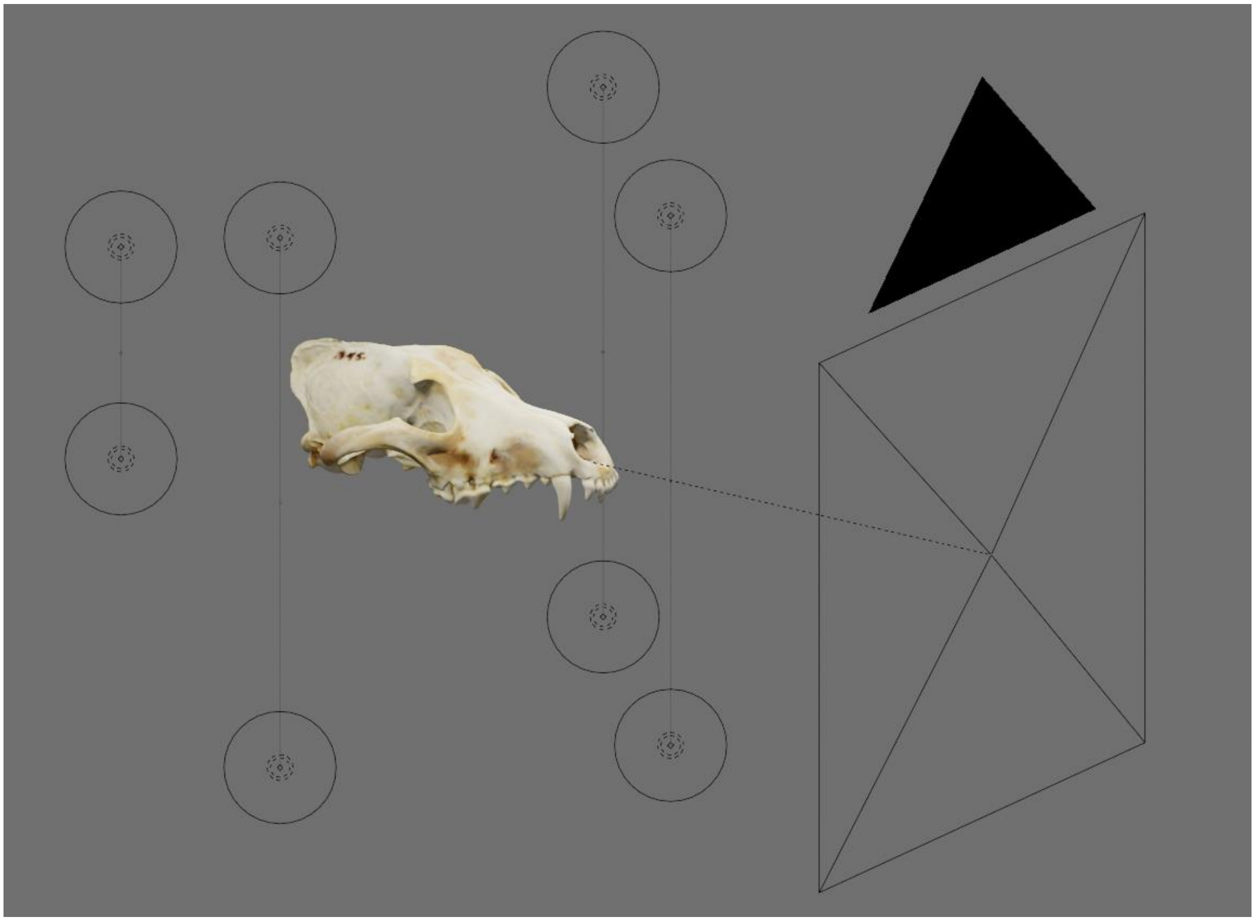

Fig S.2: The Blender environment with a wolf (*Canis lupus*) skull loaded. The background was brightened to enhance contrast of the user interface elements.

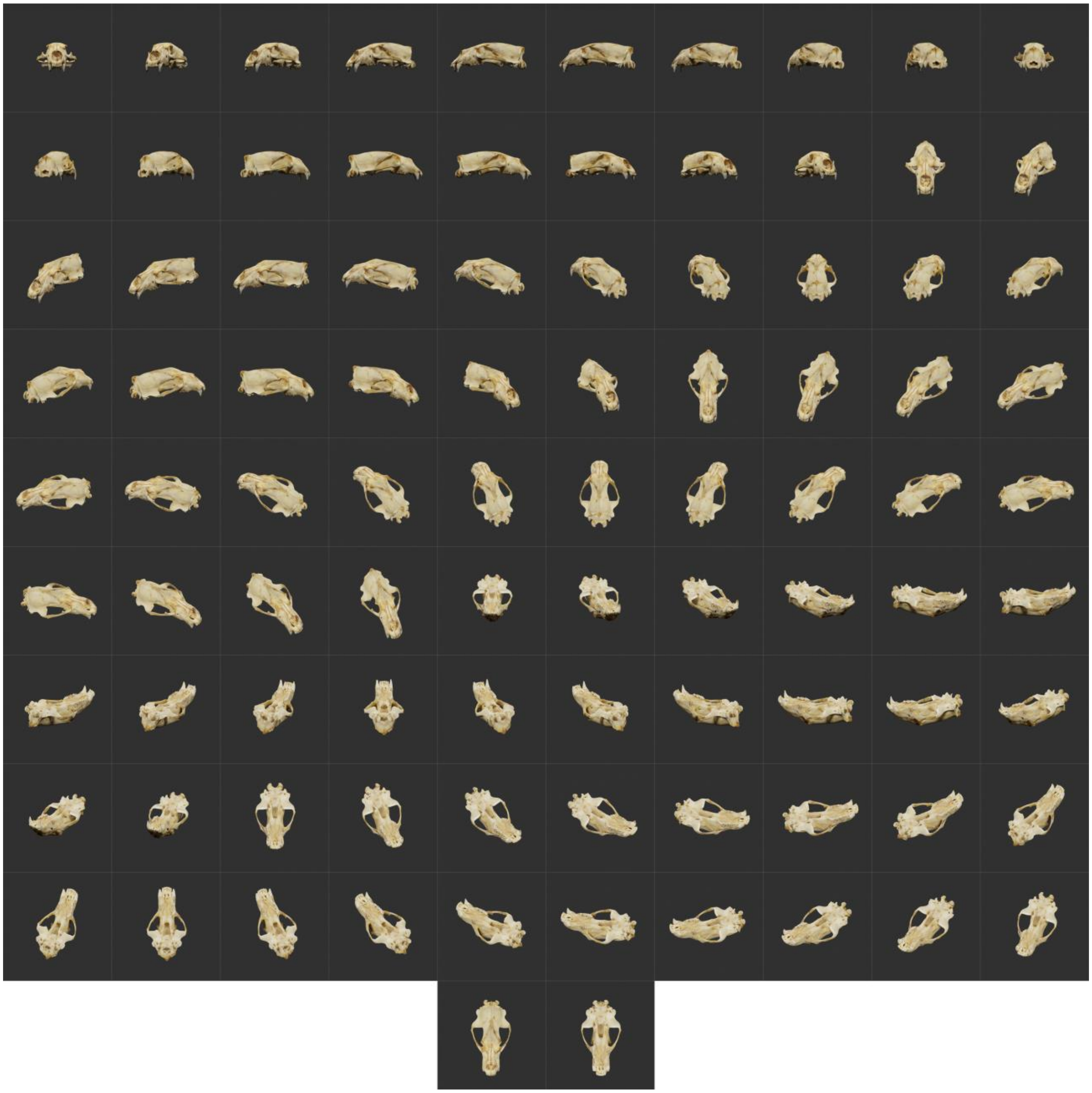

Fig S.3: A collage of synthetic images of a polar bear (*Ursus maritimus*) skull, showing all 92 imaging angles.

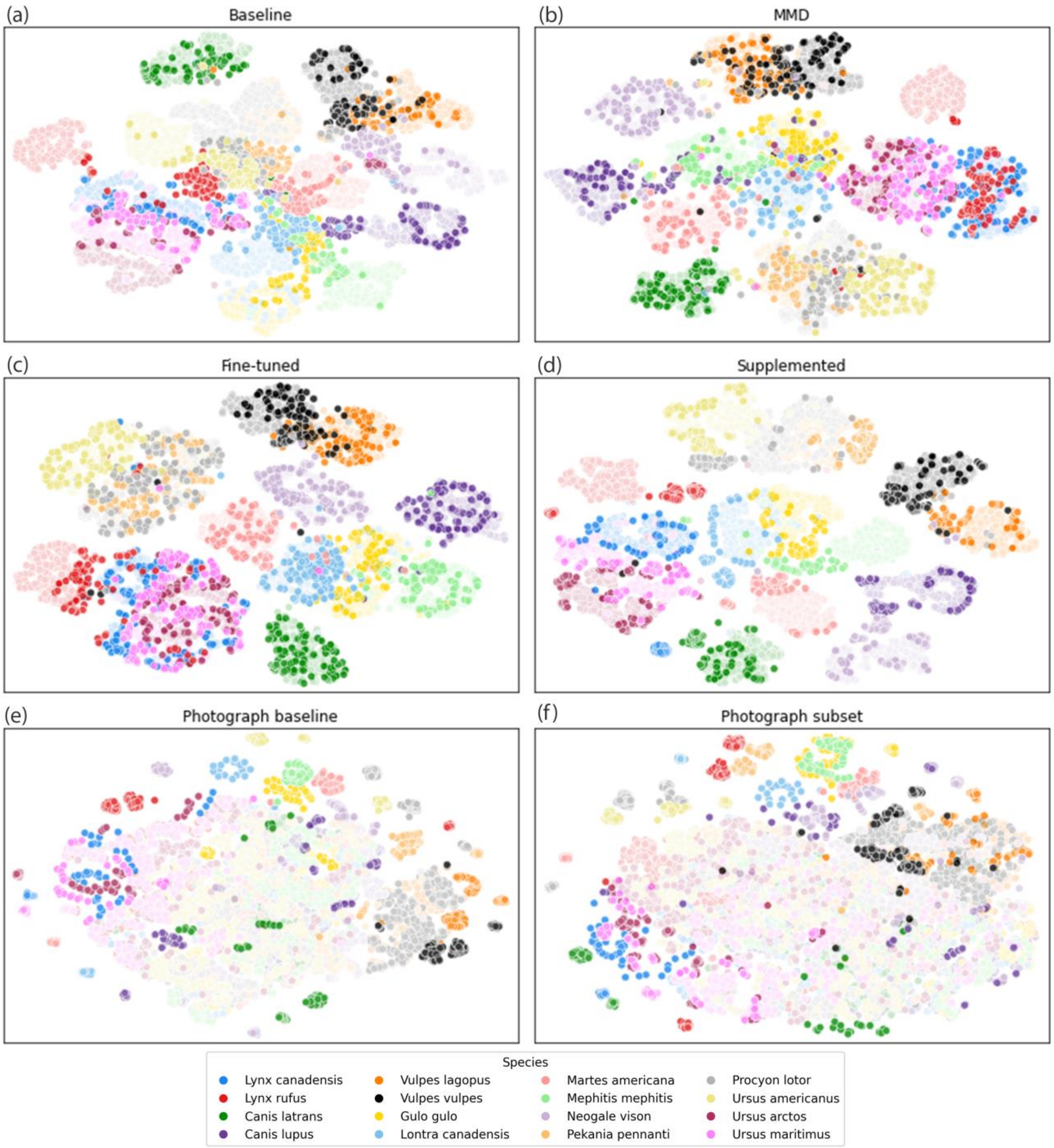


Fig S.4: Visualisation of the feature space of six skull classification models using t-SNE. Each t-SNE plot was generated from using the activations of the model's post-convolution flattened layer. Each species is represented by a unique colour. Translucent points represent synthetic images and opaque points represent photographs. All images were from the test dataset.

Tables

Table S.1: Grad-CAM heatmap scoring scheme. The second column describes the subjective criteria used to assign a Grad-CAM each score. The third column, “Examples”, shows example Grad-CAMs for each score.

| Score | Criteria | Examples |
|---|---|---|
| **3** | The heatmap appears red over the skull and blue over the background. | 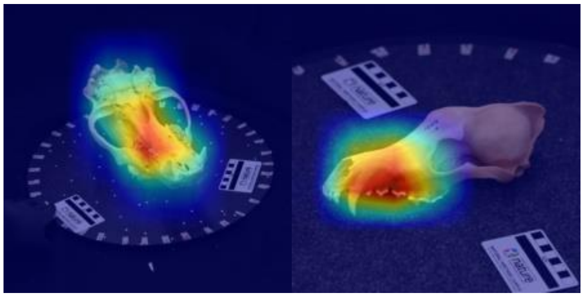 |
| **2** | **Option 1**: The heatmap appears yellow/green over the skull and blue over the background.<br>**Option 2**: The heatmap appears red over the skull and yellow/green over a small background feature (e.g., a scale card) | 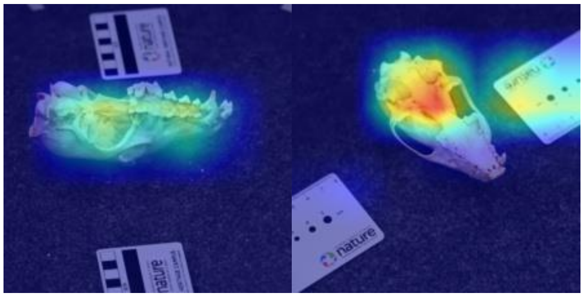 |
| **1** | **Option 1**: The heatmap shows equal activation (i.e., red/yellow/green) over the skull and a small background feature (e.g., a scale card). The rest of the background is blue.<br>**Option 2**: The heatmap appears red over the skull and green over large areas of the image’s background. | 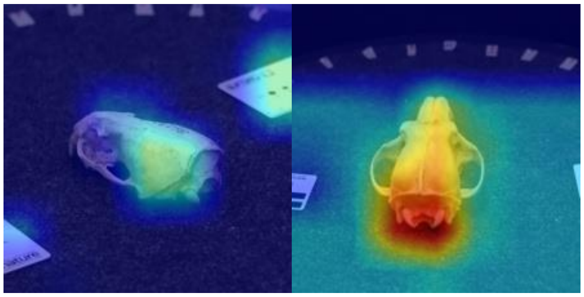 |
| **0** | **Option 1**: The heatmap shows activation (i.e., red/yellow) over large areas of the image’s background.<br>**Option 2**: The heatmap shows higher activation over a background feature than over the skull. | 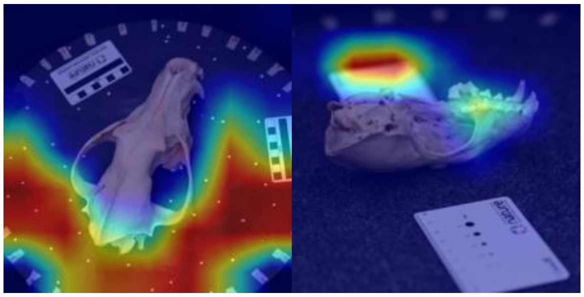 |
